# Effects of alternative blood sources on *Wolbachia* infected *Aedes aegypti* females within and across generations

**DOI:** 10.1101/414441

**Authors:** Véronique Paris, Ellen Cottingham, Perran A. Ross, Jason K. Axford, Ary A. Hoffmann

## Abstract

*Wolbachia* bacteria have been identified as a tool for reducing the transmission of arboviruses transmitted by *Aedes aegypti.* Research groups around the world are now mass rearing *Wolbachia* infected *Ae. aegypti* for deliberate release. We investigated the fitness impact of a crucial element of mass rearing: the blood meal required by female *Ae. aegypti* to lay eggs. Although *Ae. aegypti* almost exclusively feed on human blood, it is often difficult to use human blood in disease-endemic settings. When females were fed on sheep or pig blood rather than human blood, egg hatch rates decreased in all three lines tested (uninfected, or infected by *w*Mel, or *w*AlbB *Wolbachia*). This finding was particularly pronounced when fed on sheep blood, although fecundity was not affected. Some of these effects persisted after an additional generation on human blood. Attempts to keep populations on sheep and pig blood sources only partly succeeded, suggesting that strong adaptation is required to develop a stably infected line on an alternative blood source. There was a decrease in *Wolbachia* density when *Wolbachia Ae. aegypti* were fed on non-human blood sources, although density was higher again in lines kept for multiple generations on the alternate sources. These findings suggest that sheep and pig blood will entail a cost when used as alternative blood sources which needs to be taken into account when considering mass release.

## INTRODUCTION

The majority of female mosquito species require a vertebrate blood meal to obtain nutrients for successful egg development [1], with some species being explicitly adapted to humans [2,3]. Mosquito species such as *Aedes aegypti* are highly anthropophilic and are the key species involved in transmission of human pathogens such as dengue [4]. *Aedes aegypti* inhabits predominantly urban environments which currently places approximately half of the world’s population at risk of infection from dengue alone [5,6].

An emerging approach for control of dengue and other arbovirus diseases involves the release of mosquitoes infected with the endosymbiotic bacterium *Wolbachia* to reduce the transmission of the virus within human communities [7,8]. The *Wolbachia* bacterium is a maternally inherited endosymbiont that reduces the transmission of arboviruses in the *Ae. aegypti* host [9-11]. The exact mechanism that allows *Wolbachia* to induce this phenomenon is still unclear. However, factors such as a rise in host immune response, elevation in host methyltransferase to reduce viral production and suppression of key lipids required for viral replication have been identified as potentially important [12-14].

*Wolbachia* strains from other arthropod hosts have been successfully transinfected into *Ae. aegypti*, causing a range of phenotypic outcomes [15-18]. The *Wolbachia* infections *w*Mel and *w*AlbB induce cytoplasmic incompatibility (CI) in *Ae. aegypti*, resulting in embryo death when *Wolbachia*-infected males mate with uninfected females [16]. This provides a significant reproductive advantage for *Wolbachia*-infected compared to uninfected female mosquitoes. Maternal transmission and cytoplasmic incompatibility combined are mechanisms that allow *Wolbachia* infected mosquitoes to invade an uninfected population [19].

*Wolbachia* programs are initiated to suppress dengue transmission through replacing uninfected populations by infected ones or through suppressing mosquito populations using CI. Both strategies require a large number of laboratory reared mosquitoes with a high fitness under field conditions to be successful. Blood feeding is an integral part of mosquito colony maintenance. *Ae. aegypti* is highly adapted to human blood, however in many countries large volumes of human blood are difficult to obtain and subject to strict ethical and regulatory control [20] particularly given the risk of viral transfer in disease-endemic zones. Artificial blood sources are currently under development [21] and appear to act as appropriate substitutes for human blood [22] including for *w*Mel-infected mosquitoes where no effect on *Wolbachia* parameters including density, CI and virus inhibition were demonstrated [20,23]. However, issues relating to insufficient storage time [20] or reduced egg hatch rates [23] are reasons why alternative blood sources continue to be used in different locations. Where these are used, they may affect the success of releases by (1) directly decreasing the fitness of released mosquitoes, if feeding produces sub-optimal offspring that cannot compete effectively with wild mosquitoes, (2) promoting a process of genetic adaptation to non-human blood that has negative fitness consequences for released mosquitoes when feeding on human blood again, and (3) reducing *Wolbachia-*related effects like maternal transmission, CI and viral blockage [24].

Here we aimed to assess the use of non-human blood meals on *Ae. aegypti.* Previous research has shown that non-human blood sources influence the fitness of *Wolbachia*-infected mosquitoes. For instance, *w*MelPop-infected *Ae. aegypti* fed on mouse, guinea pig and chicken blood results in a sharp decrease in fecundity and egg hatch rates, in contrast to uninfected mosquitoes [25]. This raises the issue of whether there will be significant negative effects of various non-human blood sources used around the world for *Ae. aegypti* cultures.

We explore the following questions: What is the immediate impact of two common blood sources (sheep and pig) on different *Wolbachia* infection types, given that *w*Mel and *w*AlbB are now being successfully used in releases? Do the negative effects of one generation of feeding persist into the next generation if mosquitoes subsequently feed on human blood? Can large outbred populations adapt to different blood sources, and if so does this mean that performance on human blood is negatively affected following adaptation? We characterised key fitness characteristics including fecundity, egg hatch rate and larval development parameters. We also assessed *Wolbachia* density to investigate if *Ae. aegypti* fed on non-human blood sources experienced a loss or reduction of the *Wolbachia* infection.

## MATERIALS AND METHODS

### ***Wolbachia* Strains and Laboratory Conditions.**

We used *Aedes aegypti* that were uninfected or infected with the *Wolbachia* strains *w*Mel and *w*AlbB in our experiments. The *w*Mel and *w*AlbB strains are currently being reared for mass release and are described in Walker *et al.* [16], Xi *et al.* [17] and Axford, *et al.* [26]. Prior to experimentation, *Wolbachia*-infected females were crossed to uninfected Australian field caught *Ae. aegypti* males for three generations to control for genetic background effects. Each colony was maintained in a temperature-controlled insectary at 26°C ± 1°C with a 12 hr photoperiod as described by Axford *et al.* [26] and Ross, *et al.* [27]. Colonies were provided with constant access to a 10% sucrose solution and maintained in 15 cm^3^ cages (Bugdorm model #41515 Insect Rearing Cages©, Megaview Science C., Ltd., Taichung, Taiwan) and enclosed in plastic bags to maintain high humidity. Adult females were fed on a human volunteer’s arm once per generation to initiate egg laying. Colonies were maintained in two replicate cages containing approximately 100 mosquitoes each.

### Experimental Design

The study comprised two components. In the first component, the direct and intergeneration effects of blood feeding were examined. The *w*Mel, *w*AlbB and uninfected populations were randomly allocated at the larval stage into three treatments, each with 200 larvae raised with abundant food (approximately 0.5 mg/larva/day [27]). Pupae were sexed to ensure equal sex ratios. Seven-day-old females were provided with a human, pig or sheep blood meal. Combinations of blood source and *Wolbachia* infection type were initially conducted in groups of 25 females and 25 males. Blood source was scored for fecundity, egg hatch rate and *Wolbachia* density. To test for persistent effects of blood feeding, F1 offspring from females fed on different blood sources were then fed on human blood and re-tested for fecundity, egg hatch rate and *Wolbachia* density.

In the second component, we determined if adaptation to non-human blood sources was possible and costs associated with this process. *Aedes aegypti* colonies that were uninfected or carrying the *w*AlbB or *w*Mel infections were maintained on pig, sheep or human blood for four generations. We attempted to maintain a large population size in these experiments to avoid deleterious inbreeding effects that can be substantial in *Ae. aegypti* by turning over 200+ adults each generation, but terminated lines once these fell below 20 because inbreeding effects then make line comparisons difficult [28]. This experiment was run on unreplicated lines because of resource limitations. Unfortunately, some lines were lost in this experiment presumably due to the negative effects of maintaining mosquitoes on alternate blood sources (see below) which may have been cumulative across time. The *w*Mel and *w*AlbB lines maintained on sheep were lost at the second and third generation respectively. The *w*Mel line maintained on pig blood was lost at the F4 stage when <20 larvae developed. The remaining lines were scored for egg hatch rate and development time after four generations (F4). *Wolbachia* density was also scored at the start of this experiment in F1 offspring after parents had been held for a generation on different blood sources (ie., before any lines were lost).

We continued the surviving lines on their respective blood sources for an additional six generations (i.e. F10, 10 generations in total), with no further lines lost because large population sizes had developed by this stage and these were easily maintained. To test for any deleterious effects of the selection process, mosquitoes from these lines were fed on human blood and retested for fecundity, egg hatch rate and *Wolbachia* density.

### Blood feeding

Pig and sheep blood were considered for their potential as an alternative blood source to human blood because of their ready availability in some countries where releases are planned or being undertaken. Fresh sheep and pig blood were collected from JBS Australia© (Brooklyn, Australia) and Diamond Valley Pork© (Laverton North, Australia), respectively, into 10 mL Lithium Heparin Vacuette evacuated blood collection tubes (Greiner Bio-One International, Austria) and stored at 4°C. Before blood collection, animals were kept for two weeks without access to antibiotic containing food and water sources to ensure that the blood was mostly free of antibiotics. Human blood was sourced from the Red Cross (Agreement # 16-10VIC-02). Colonies were tested after being maintained on their respective blood sources or being transferred back to human blood. The *w*Mel groups fed on sheep and pig blood and the *w*AlbB group fed on sheep blood were not tested later because population sizes declined, resulting in insufficient egg numbers to allow the populations to persist without severe inbreeding effects being anticipated [28].

Mosquitoes were fed via Hemotek© membrane feeders (Discovery Workshops, Accrington, UK) [29]. To prepare the blood-filled Hemotek© disc, a 6 cm x 6 cm square of collagen membrane (Discovery Worskshops, Accrington, UK) was cut out and placed over the metal reservoir. The membrane was then fastened with an O ring. Approximately 6 mL of blood was pipetted into the metal reservoir which was plugged with two nylon stoppers. The blood-filled reservoir was screwed onto the heat-supplying feeder, calibrated to 37 °C (human body temperature). The completed feeder was placed on top of the mosquito colony cage. Colonies were given approximately 20-25 minutes to feed, or until all females showing an interest in feeding by probing the blood delivery site with their proboscis had taken a blood meal (indicated by an engorged abdomen). Blood meal reservoirs were restricted to *Wolbachia* infection types, i.e. there was no sharing of blood reservoirs between infection types to avoid the unlikely event of cross-contamination.

### Fecundity and Egg Hatch Rate

Fecundity was measured by isolating engorged females. For the first component looking at parental effects following feeding on different blood sources or subsequent feeding on human blood, five to six females per replicate (4 cages per *Wolbachi*a infection type/blood source group) were taken (for a total of 20-24 females for each *Wolbachia* infection/blood source group). For the second component (following selection on different blood sources), 2-3 females per replicate were taken in the F4 generation from each of three cages (for a total of 5-9 females per *Wolbachia* infection/blood source group). More females were available at the F10 generation for testing when 16-24 females were used per blood source.

For the assays, engorged females were isolated in 70 mL cups filled halfway with reverse osmosis (RO) water and lined with sandpaper (Norton^®^ Master Painters P80, Saint-Gobain Abrasives Pty. Ltd., Thomastown, Victoria, Australia) as an oviposition substrate. The top of the cup was covered with mesh to prevent mosquitoes escaping. Females were allowed one week to lay eggs before being stored in 70 % ethanol and sandpaper strips were collected and partially dried. Three days after collection, eggs were hatched in plastic trays filled with 500 mL of RO water. Two days later, fecundity and egg hatch rates were calculated by counting the number of hatched and unhatched eggs. An egg was recorded as ‘hatched’ if the egg cap was detached.

### Development time

After lines had been cultured for several generations on different blood sources as part of the second component, we noticed that there was a delay in larval development. We quantified development time at the F4 generation as the duration taken from hatching to adult emergence. Larvae were selected at random using a pipette and placed into trays filled with 150 mL of RO water where they were given access to TetraMin^®^ fish food (Tetra, Melle, Germany) *ad libitum* (approximately 0.5 mg/larvae/day). Trays of larvae were maintained in incubators at 26 °C ± 1 °C. Trays were rotated daily to account for any variation in temperature or light within the incubators. Trays were checked twice daily where the number and sex of emerging adults was recorded. Development time was scored on 4-22 individuals of each sex per line from a particular blood source.

#### **DNA Extraction and *Wolbachia* Screening**

We extracted genomic DNA using Chelex^®^ 100 Resin (Bio-Rad Laboratories, Hercules, CA). Adults were removed from cages for extraction one week after blood feeding to ensure that eggs had been laid. Extraction was conducted using 250 µl of 5 % Chelex^®^ and 3 µl of Proteinase K (20 mg/ mL) (Roche Diagnostics Australia Pty. Ltd., Castle Hill New South Wales, Australia) solution. Individual samples were ground using two 3 mm glass beads per sample in a mixer mill (Retsch^®^ GmbH, Haan, Germany). Samples were then incubated for 60 minutes at 65 °C and 10 minutes at 90 °C. Samples were diluted by 1:10 then stored at - 20 °C.

We used a LightCycler^®^ 480 to confirm *Wolbachia* infection status and estimate *Wolbachia* density. Three primer sets were used to amplify markers specific to the *Aedes* genus, differentiate *Ae. aegypti* from other *Aedes* species and amplify markers specific to the *Wolbachia* infections (*w*Mel or *w*AlbB) as described by Lee *et al.* [30] and Axford *et al.* [26], where forward and reverse primer sequences can be found. Comparisons between these three sets of markers was based on crossing point (Cp) values and temperature melt curve (Tm) analysis. Three technical replicates were completed for each mosquito; differences in crossing point between the two markers were averaged to obtain an estimate of *Wolbachia* density (once adjusted for the *Aedes* control gene Cp value). Detailed analysis of the qPCR protocol is provided by Yeap *et al.* [31] and Lee *et al.* [30].

We estimated the density of the *w*Mel and *w*AlbB infections from each generation and blood source by comparing *Ae. aegypti* and *Wolbachia* Cp values. The *Wolbachia* Cp value (Cp2) was subtracted from the *Ae. aegypti* Cp value (Cp1) as a relative estimate of *Wolbachia* density. If we assume that the efficiency of the reaction is perfect (i.e., R = 2^-ΔΔCp^), the difference in Cp values between samples can give an estimate of the relative concentration of *Wolbachia* DNA relative to mosquito DNA when computed as 2^(Cp1–Cp2)^. While Cp values do not give definitive quantification, they provide a relative guide as to the concentration of a *Wolbachia* infection when compared to the host. However in other work we have found a good agreement between relative *Wolbachia* densities established through Cp values with those obtained via a Droplet Digital system that directly measures absolute copy number [32].

#### Statistical Analysis

Statistical analysis was completed using SPSS V21.0 for Windows (SPSS Inc, Chicago, IL) and R (0.99.902)[33]. All data sets were tested for normality using Shapiro-Wilks tests. Data sets were analysed through general linear models with and without transformation, although in no case did the transformation influence the conclusions from the analysis. For the results, analyses on egg hatch rates are presented after these had been angular transformed, while analyses of fecundity data are presented after these were ln(x+1) transformed, however both variables are displayed as box plots on the untransformed data. Tukey’s b posthoc test was undertaken for pairwise comparison between blood sources and *Wolbachia* infection type where appropriate. Density data (*Wolbachia* density relative to *Aedes* controls) were also analysed through general linear models.

## RESULTS

### Immediate and F1 effects (component 1)

#### Fitness effects

The fecundity of the three infection types (uninfected, *w*Mel and *w*AlbB) was not affected by the blood source (General Linear Model: F_(2, 279)_ = 2.129, P = 0.121). While there was no interaction between *Wolbachia* infection type and blood source (F_(4,279)_ = 0.836, P = 0.503), there was an effect of the infection type overall (F_(2,279)_ = 3.267, P = 0.040) reflecting the fact that the *w*AlbB strain had somewhat higher fecundity than *w*Mel in this experiment (Figure 1A).

**Figure 1.**
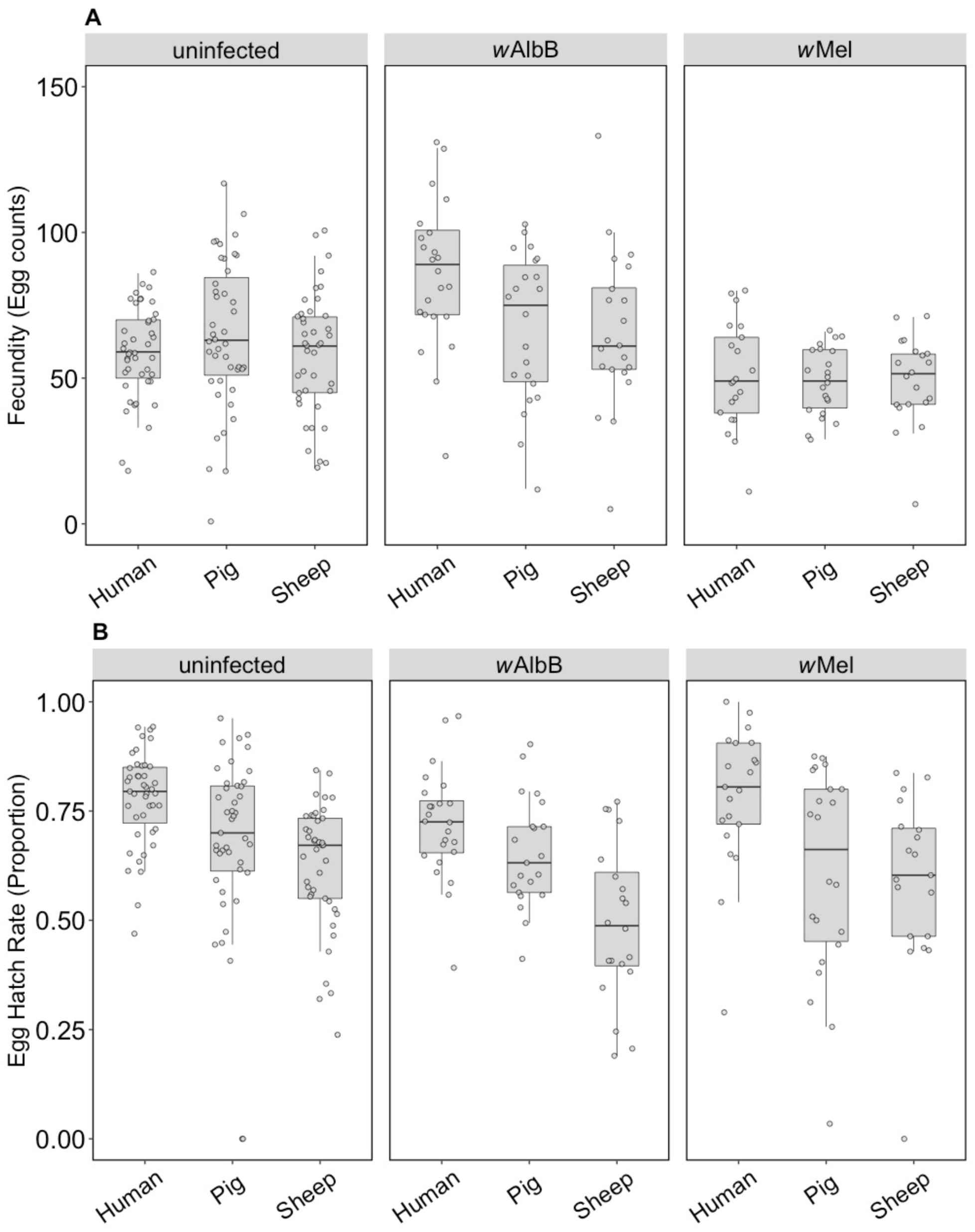
Comparison of Fecundity and Egg Hatch Rates across infection types (uninfected, wMel, wAlbB) and blood sources (human, pig, sheep). (A) Fecundity (egg counts) (B) Egg Hatch Rate (Proportion). Each dot embodies the egg count and egg hatch rate of an individual female. The boxplots represent the first to the third quartiles around the median (horizontal grey line) and the vertical bars the 1.5 interquartile of the lower and upper quartiles. Around 24 females per blood source and generation were used in this analysis.

In contrast to fecundity, we found a significant effect of the blood source (General Linear Model: F_(2,246)_ = 29.32, P < 0.001) on egg hatch rates. All *Wolbachia* infection types (uninfected, *w*Mel and *w*AlbB) fed on non-human blood sources showed a significant decrease in egg hatch rate compared to mosquitoes fed on human blood (post hoc tests, all p < 0.001) (Figure 1B). There was also an effect of *Wolbachia* infection type (F_(2,246)_ = 4.795, P = 0.009) but no interaction between infection type and blood source (F_(4, 246)_ = 0.195). Posthoc tests indicated that *w*AlbB egg hatch rate was lower than that of the uninfected group, while *w*Mel overlapped with both (Figure 1B). The total fertility of a female fed on sheep blood as measured by the number of larvae was 36.3% lower than for a female fed on human blood, while pig blood feeding reduced this by 16.5%.

The F1 generation of parents fed on non-human blood sources showed persistent effects of parental blood feeding despite F1 exposure to human blood. There was no effect of parental blood source (F_(2,279)_=0.399, P = 0.671), infection type (F_(2, 279)_ = 0.032, P = 0.968) or an interaction effect (F_(4,279)_=0.083, P = 0.987) on (ln (x+1)) fecundity (Figure 2A). However, (angular transformed) egg hatch rates were affected by parental blood source (F_(2,247)_=5.476, P = 0.005), with reduced egg hatch rates when parents were fed on pig or sheep blood (Figure 2B). There were no effects of the infection type (F_(2,247)_ = 0.238, P = 0.789) or interaction between infection type and blood source (F_(4,247)_=1.063, P = 0.375) on egg hatch rates. The fertility of a female whose mother had fed on sheep blood as measured by the number of viable eggs was 16.8% lower than for a female fed both generations on human blood, while pig blood feeding by a mother reduced fertility by 7.9%.

**Figure 2.**
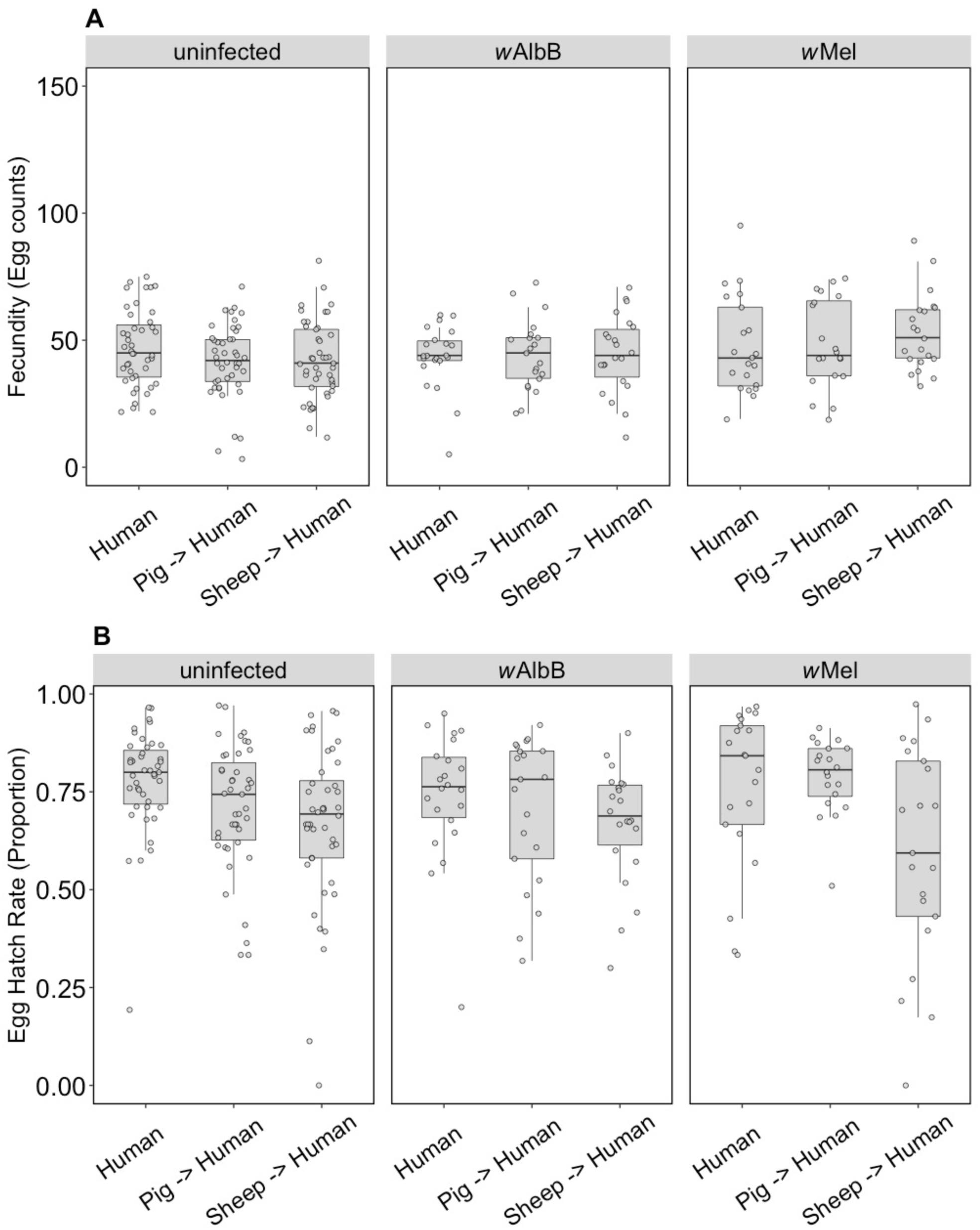
Comparison of Fecundity and Egg Hatch Rates across infection types (uninfected, wMel, wAlbB) when females were fed on human blood but parents were fed on different blood sources (human, pig, sheep). (A) Fecundity (Egg counts) (B) Egg Hatch Rate (Proportion). Each dot embodies the egg count and egg hatch rate of an individual female. The boxplots represent the first to the third quartiles around the median (horizontal grey line) and the vertical bars the 1.5 interquartile of the lower and upper quartiles. Around 24 females per blood source and generation were used in this analysis.

##### Wolbachia *density*

We compared the *Wolbachia* density of parents fed on different blood sources and the density of F1s fed on human blood. For *w*AlbB in the parents, there was no effect of blood source on *Wolbachia* density (F_(2,33)_ = 1.138, P = 0.333) but there was for *w*Mel (F_(2,33)_ = 5.079, P = 0.012), with density being lower when females were fed on pig and sheep blood (Figure 3A). The *w*AlbB density when F1s were fed on human blood was affected by parental blood source (F_(2,33)_ = 9.223, P = 0.001); density was lower when parents were fed on both pig and sheep blood (Figure 3B), with both treatments differing in posthoc tests from the human blood treatment. There was also a difference in density between the F1s for *w*Mel (F_(2,33)_ = 8.140, P = 0.001), with a reduced density when parents were fed on pig and sheep blood even though F1s had been fed on human blood (Figure 3B).

**Figure 3.**
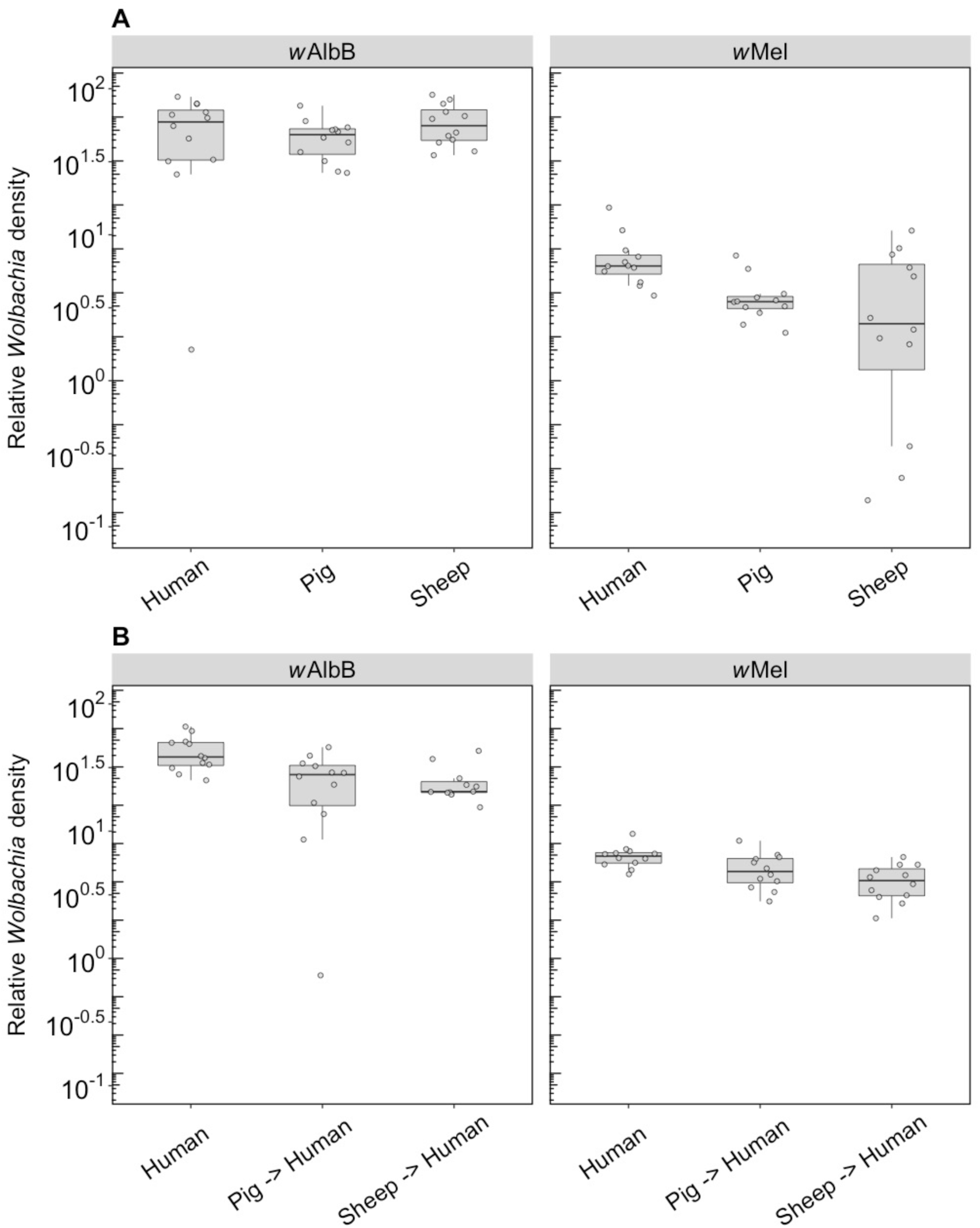
Comparison of the Relative Wolbachia density of the infections types (wMel, wAlbB) across different blood sources (human, pig, sheep). (A) Relative Wolbachia density of parents fed on different blood sources (B) Relative Wolbachia density of F1s after being re-introduced to human blood. Each dot represents the density of an individual mosquito. The boxplots represent the first to the third quartiles around the median (horizontal grey line) and the vertical bars the 1.5 interquartile of the lower and upper quartiles. Around 12 individuals per blood sources were used in the F1 and F2 generation.

### Long term effects (Component 2)

#### Fitness effects (F4)

We compared the fecundity and egg hatch rates of lines kept on the three blood sources for four generations. Note that the number of females in these tests (5-9) was lower per treatment group than in the earlier trials, which was a reflection of the difficulty we had in maintaining mosquitoes on the different blood sources. All infection types performed well on human blood and there were no differences in fecundity (F_(2,16)_ = 2.547, P = 0.110) (Figure 4A) or egg hatch rate (F_(2,16)_ = 0.757, P = 0.485) (Figure 4B). For groups maintained on pig blood, both fecundity (F_(1,16)_ = 5.414, P = 0.033) and egg hatch rate (F_(1,16)_ = 32.652, P < 0.001) were lower in *w*AlbB mosquitoes compared to uninfected mosquitoes (Figure 4A,B).

**Figure 4.**
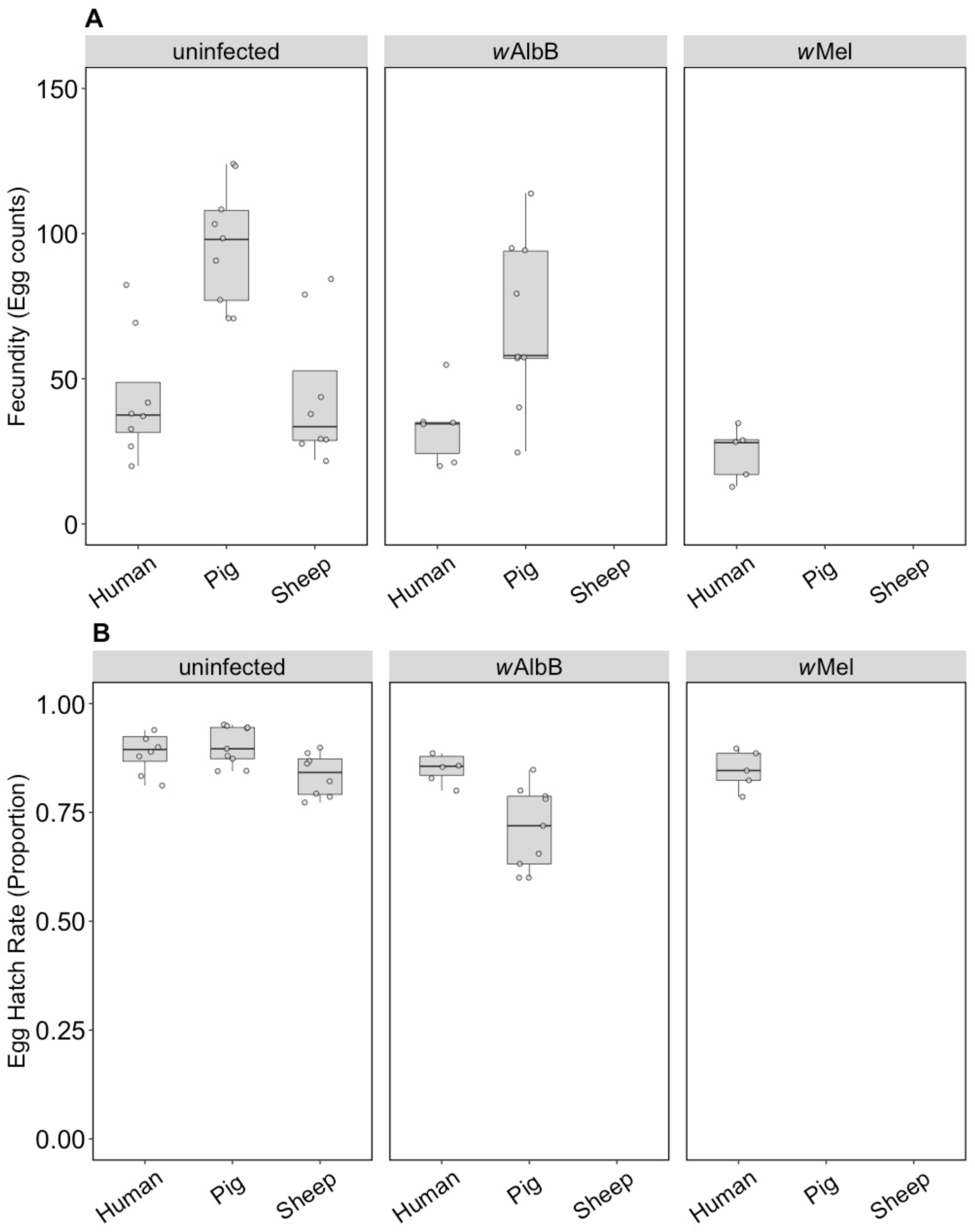
Comparison of Fecundity and Egg Hatch Rates across infection types (uninfected, wMel, wAlbB) and different blood sources (human, pig, sheep) of F4 selected groups. (A) Fecundity (Egg counts) (B) Egg Hatch Rate (Proportion). Each dot embodies the egg count and egg hatch rate of an individual female with 5-9 females tested per treatment. The boxplots represent the first to the third quartiles around the median (horizontal grey line) and the vertical bars the 1.5 interquartile of the lower and upper quartiles.

For the uninfected group maintained on pig or sheep blood, fecundity differed between blood sources (F_(2,22)_ = 13.14, P < 0.001) and was higher when mosquitoes were fed on pig blood compared to human or sheep (Figure 4A), though egg hatch rates did not differ significantly (F_(2,22)_ = 3.21, P = 0.060) (Figure 4B). This resulted in a 127% increase in viable eggs when mosquitoes fed on pig *versus* human blood and a small reduction of 4.5% for sheep. Likewise, *w*AlbB exhibited significantly higher fecundity on pig compared to human blood (F_(1,13)_ = 8.87, P = 0.011) but egg hatch rates were lower (F_(1,13)_ = 10.05, P = 0.007), however there was still a 69.9% increase in larval number on pig blood. These results contrast to those obtained in the parental generation, suggesting that there is blood source adaptation in the lines.

For the uninfected group where all three blood sources could be compared, there was a significant effect of blood source on development time (F_(2,16)_ = 19.46, P < 0.001). Larval development of the uninfected group held on human blood was consistently faster than that of the pig group and to a lesser extent the sheep group, regardless of sex (Figure 5A,B). There was no significant effect of sex (F_(1, 16)_ = 3.832, P= 0.068) or any interaction between sex and blood source (F_(2,16)_ = 0.092, P = 0.912). For the *w*AlbB lines, there was no significant effect of blood source on development time overall (F_(1, 12)_ = 2.757, P = 0.123) (Figure 5A,B).

**Figure 5.**
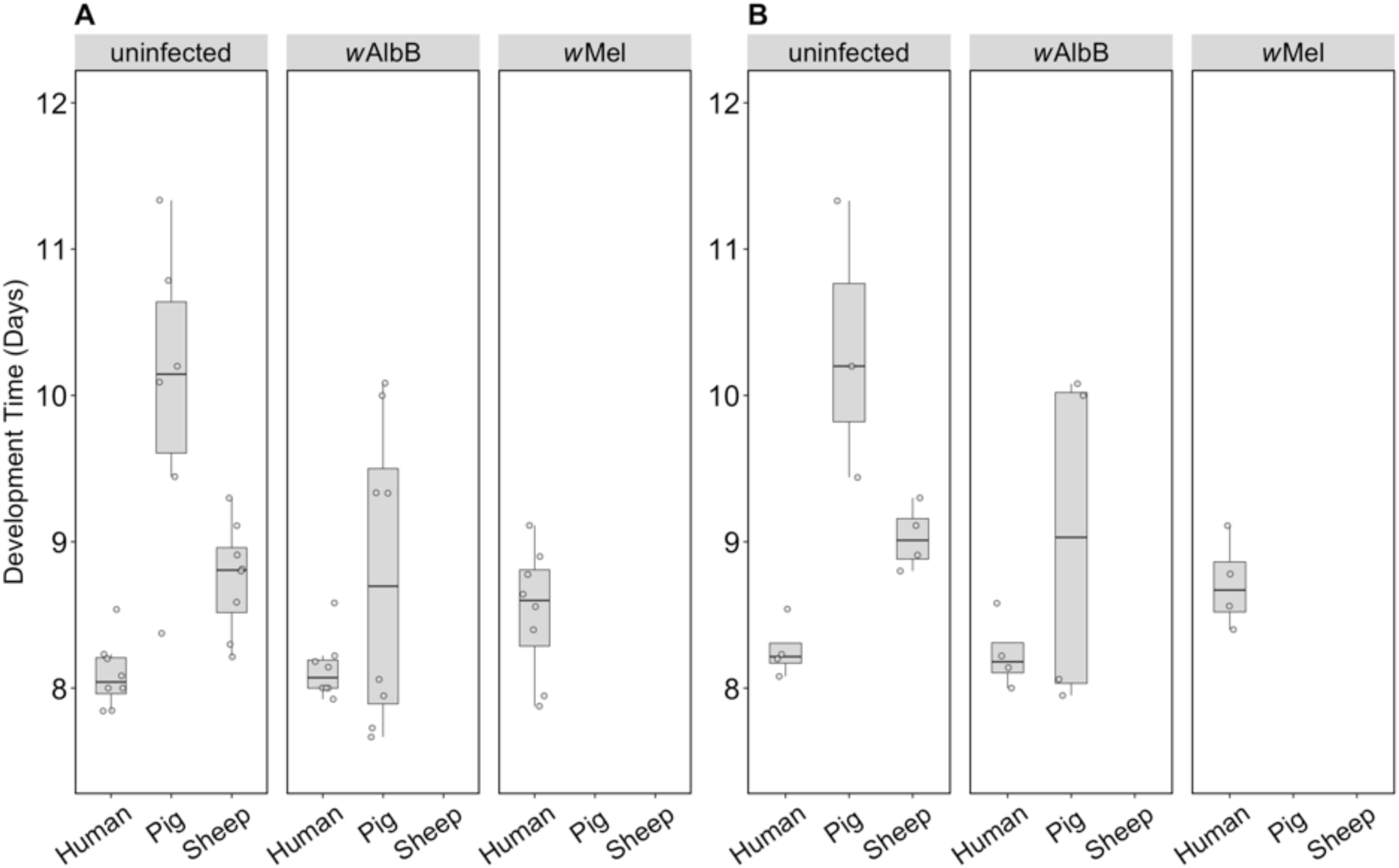
Comparison of Development Time across infection types (uninfected, wMel, wAlbB) and different blood sources (human, pig, sheep). The development time in days, calculated as the number of days from being hatched (day 0) to emerging as adults. (A) Males (B) Females. Individual data points indicate the averaged development time of mosquitoes in each replicate based on 4-22 individuals. The boxplots represent the first to the third quartiles around the median (horizontal grey line) and the vertical bars the 1.5 interquartile of the lower and upper quartiles.

#### Wolbachia *density (F1)*

F1s from parents fed on either pig or sheep blood showed a two to four-fold drop in density compared to parents that fed on this blood (c.f. Figure 6 with Figure 3). Differences in density were highly significant depending on blood source. For *w*AlbB, there was a highly significant difference (F_(2,26)_ = 53.98, p<0.001) with a 5-fold density difference between parents fed pig and human blood and a 3.5-fold difference between parents fed sheep and human blood. In contrast, exposure to different blood in parental mosquitoes resulted in similar densities of this infection across groups (Figure 3). For *w*Mel there was also a significant difference among blood sources (F_(2,24)_ = 19.08, P < 0.001). A decrease in F1s from parents with pig and sheep blood feeding had previously been evident from direct feeding for this infection type (Figure 3) although in the F1s the density on pig was lower than on sheep whereas this was reversed under direct exposure.

**Figure 6.**
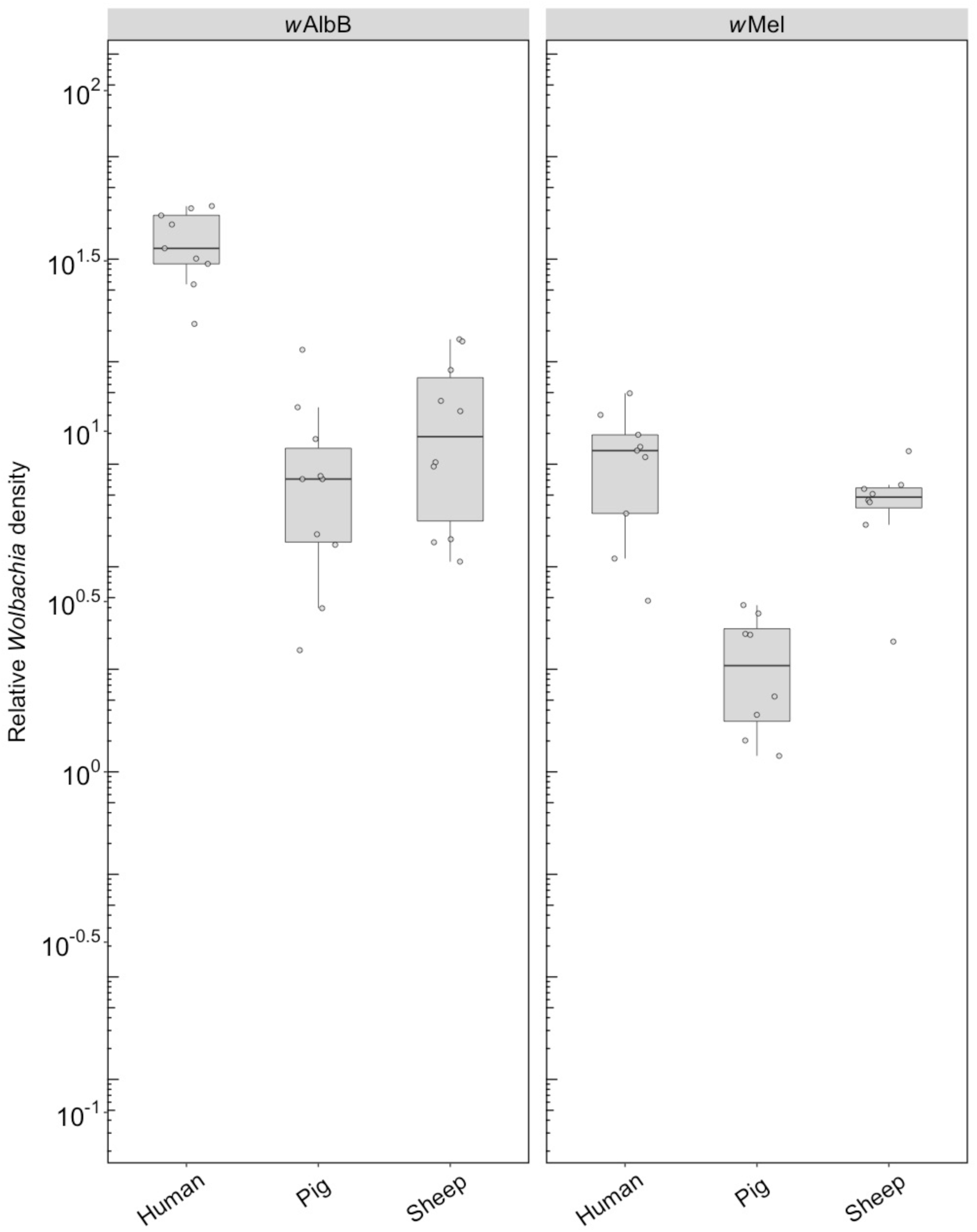
Comparison of the Relative Wolbachia density of the infections types (wMel, wAlbB) across different blood sources (human, pig, sheep) in the F1s that represent the offspring of parents exposed to their respective blood source. Each dot represents the density of an individual mosquito. The boxplots represent the first to the third quartiles around the median (horizontal grey line) and the vertical bars the 1.5 interquartile of the lower and upper quartiles. Between 8 and 10 F1s per blood source were scored.

#### Fitness effects (F10, after feeding on human blood)

The groups placed on the different blood sources were successfully maintained until generation 10, consistent with an ability of these populations to adapt to blood sources once they are established. We were then interested in testing whether there was any cost in terms of maintaining fitness when groups were switched back to human blood. For the uninfected line, there was no change in fecundity associated with blood feeding selection (F_(2, 57)_ = 0.523, P = 0.596) (Figure 7A) but there was an effect on egg hatch rate (F_(2, 56)_ = 7.16, P = 0.002) due only to sheep compared to the other two blood sources (Figure 7B) in a posthoc analysis. For the *w*AlbB infection type, there was no change in fecundity associated with blood feeding selection on pig *versus* human (F_(1, 35)_ = 0.150, P = 0.700) and egg hatch rate was also similar (F_(1, 35)_ = 1.431, P = 0.240) (Figure 7A,B).

**Figure 7.**
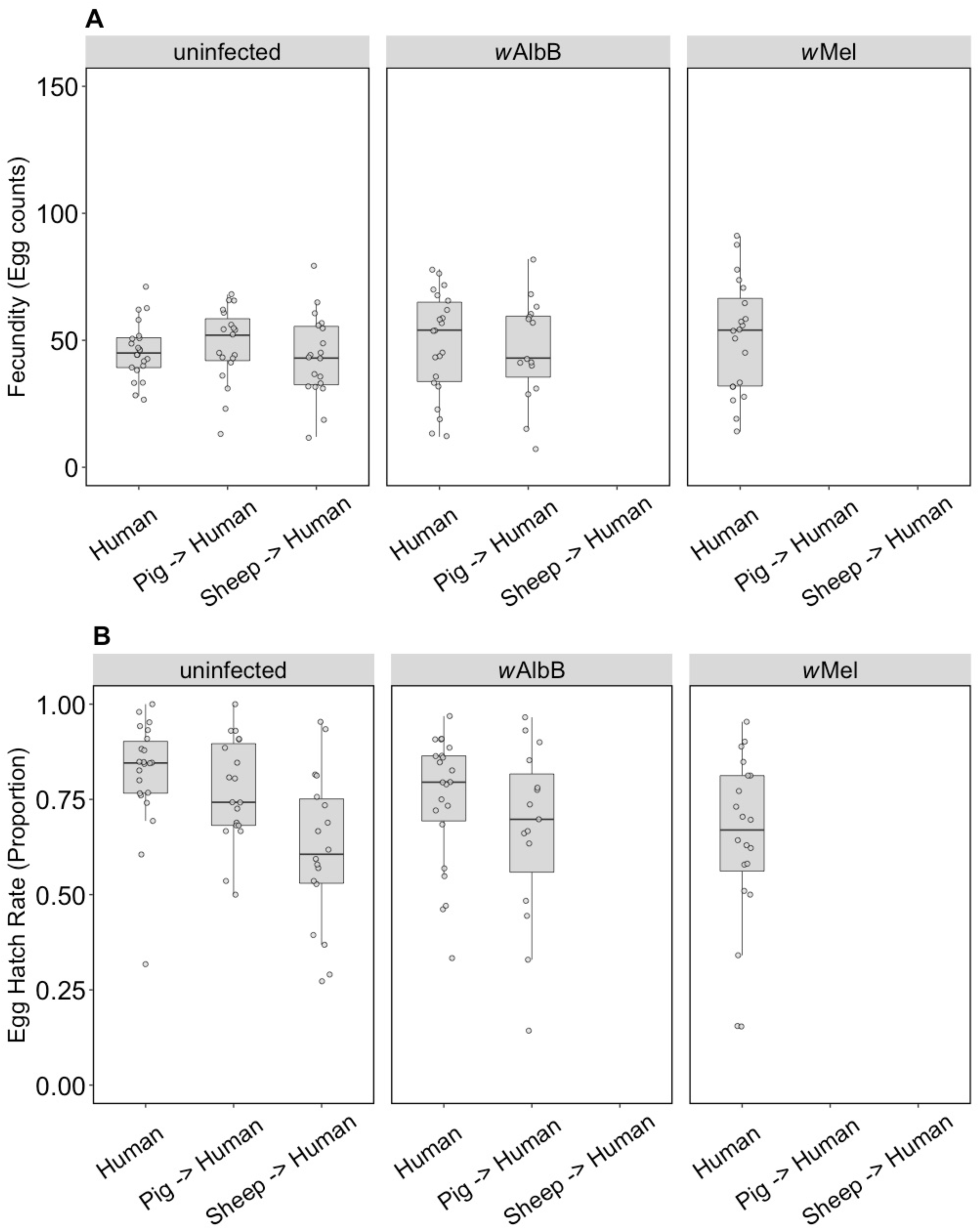
Comparison of Fecundity and Egg Hatch Rates across infection types (uninfected, wMel, wAlbB) when females were fed on human blood after colonies were maintained on different blood sources for 10 generations (human, pig, sheep). Some of these groups lost their infection during maintenance, therefore no comparison is possible for wAlbB and wMel-infected groups fed on sheep blood. There is also no wMel-infected group maintained on pig blood. (A) Fecundity (Egg counts) (B) Egg Hatch Rate (Proportion). Each dot embodies the egg count and egg hatch rate of an individual female. The boxplots represent the first to the third quartiles around the median (horizontal grey line) and the vertical bars the 1.5 interquartile of the lower and upper quartiles. Around 15-24 females per blood source and generation were used in this analysis.

#### Wolbachia *density (F10, after feeding on human blood)*

**Figure 8.**
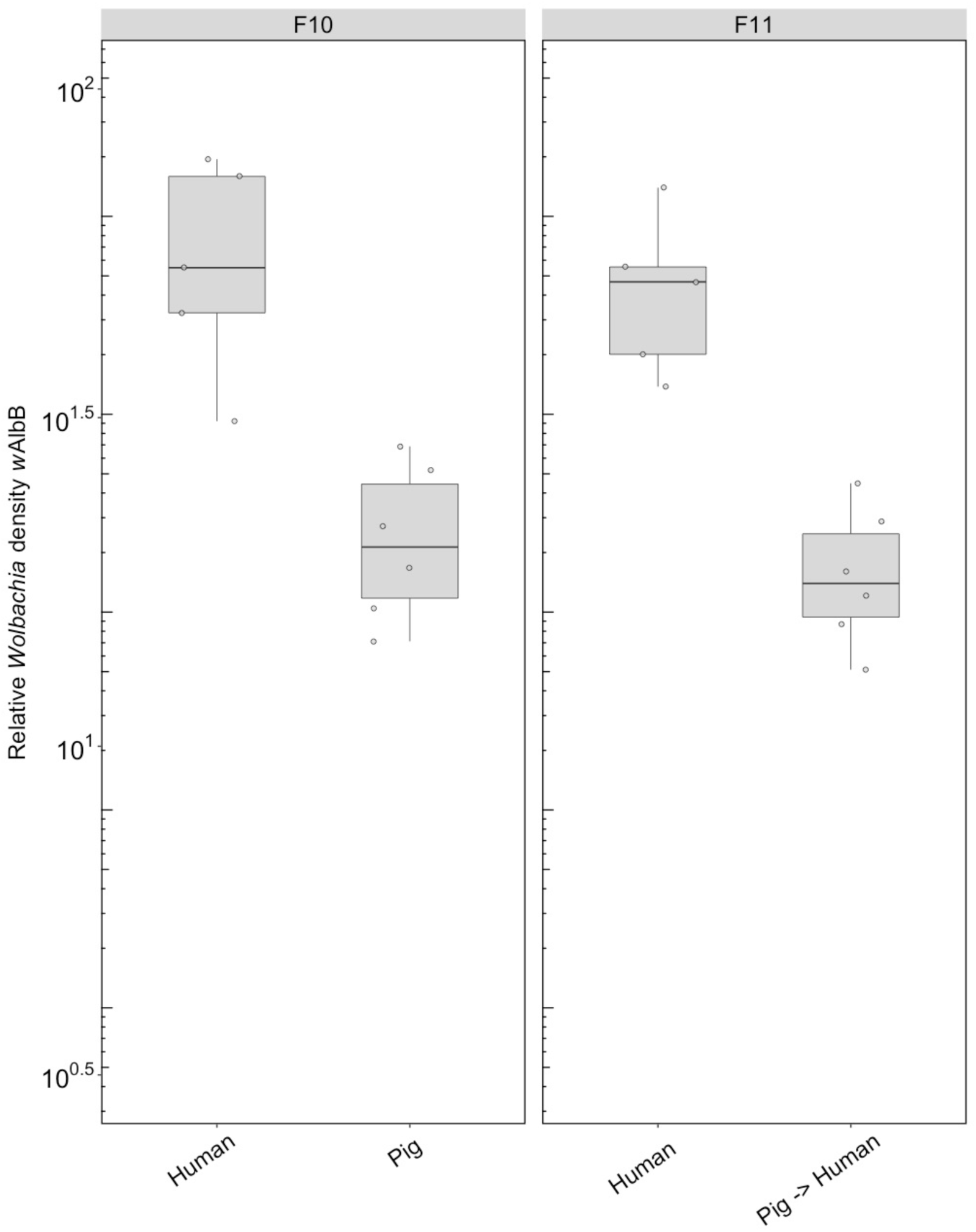
Comparison of the Relative Wolbachia density of the wAlbB infections across the different blood sources (human, pig) after 10 generations of maintenance. Some of these groups lost their infection during maintenance, therefore no comparison is possible for wAlbB and wMel infected groups fed on sheep blood. There is also no wMel infected group selected on pig blood. Each dot represents the density of an individual mosquito. The boxplots represent the first to the third quartiles around the median (horizontal grey line) and the vertical bars the 1.5 interquartile of the lower and upper quartiles. Around 6 individuals per blood source were used in the analysis.

## DISCUSSION

This study indicates that key reproductive traits in *Ae. aegypti* can be lowered by a sheep or pig blood meal *versus* a human blood meal. While many mosquito species utilise a wide variety of blood sources to facilitate the egg laying process, *Ae. aegypti* appears dependent on human blood for this purpose [24,34]. Both groups fed on non-human blood showed a significant decrease in egg hatch rates compared to mosquitoes fed on human blood, regardless of their *Wolbachia* infection. Moreover, there appears to be a carry-over effect when the mosquitoes feed on human blood in the next generation. When considering that laboratory reared mosquitoes are to be released into field environments with access to human hosts, we tested the effect of a single human blood meal of groups previously fed on human, pig and sheep blood to investigate if a carry-over effect of fitness effects can be observed. After reintroducing the F1 generation of each blood source (human, pig, sheep) to human blood, an increase in egg hatch rates was observed for the non-human parental exposure treatments. The egg hatch rates of the group previously fed on sheep blood remained significantly lower compared to the groups exclusively fed on human blood. Considering these results with respect to vector control release programs, the use of a non-human blood source (i.e. sheep blood) could not only lead to a reduced fitness of released mosquitoes but may also reduce the reproductive success of the next generation.

Feeding on non-human blood sources also seems to limit the potential for optimal *Wolbachia-* infected *Ae. aegypti* colony maintenance, consistent with previous findings [24,35,36]. We were unable to sustain the *w*Mel groups on pig and sheep blood past the F3 and F2 respectively due to low numbers of larvae. The *w*AlbB group maintained on sheep blood was also lost at the F3 stage. While larval population numbers of the *w*AlbB sheep group were high (two replicates, each with approximately 200 larvae), most of this group was found dead as 3^rd^-4^th^ instar larvae. We are unclear about the reasons for this mortality in the groups, but perhaps *Wolbachia* generates additional costs on larvae when these are insufficiently provisioned after feeding on alternate blood sources. It is possible that in the absence of access to nutrients supplied by the blood meal, the *Wolbachia* infection may utilize key resources from the host. This is supported by the finding that *Ae. aegypti* cholesterol levels are reduced by 15-25 % when carrying a *Wolbachia* infection compared to the absence of a *Wolbachia* infection [36]. In contrast, all the infected groups could be easily maintained on human blood for multiple generations. Other factors like inbreeding may also be involved if a low proportion of females in each population fed successfully across the generations, resulting in bottlenecks [28].

When considering long-term fitness effects *Ae. aegypti* may experience if fed on non-human blood sources for several generations, we found a selection response in the uninfected controls, in that egg hatch rates on alternate blood sources were relatively high by F4. There was also an apparent increase in fecundity in the population held on pig blood. However, egg hatch rates still showed a cost. Moreover, the egg hatch rate effect persisted when mosquitoes were fed human blood again even after 10 generations of maintenance, suggesting a long-term fitness cost.

For the effects of blood meal on *Wolbachia* density, we found different patterns for the two infection types (*w*Mel & *w*AlbB) as well as differences between short- and long-term effects. After one generation of either pig or sheep blood, *w*Mel showed a significant drop in density, whereas the density of *w*AlbB remained almost the same. Both infection types did not show an increase in density after being reintroduced to human blood. Considering the long-term effect on the *Wolbachia* infection, we found a 4-fold drop in density exhibited by *w*AlbB and *w*Mel when fed on either sheep or pig blood. We are unsure as to why the *Wolbachia* infection density is so negatively affected by alternative blood sources. However, it has been documented previously that endosymbionts often induce a reduction in host cholesterol levels [37,38]. It has been suggested this is due to competition between the host (mosquito) and the *Wolbachia* endosymbiont for key nutrients supplied by the blood meal, yet there is also a high degree of disparity between the cholesterol and amino acids between human and other animal blood sources [36]. When considering *w*MelPop and *w*Mel-infected *Ae. aegypti,* Caragata *et al.* [36] identified a rapid reduction in fecundity and viability (egg hatch rate) of the eggs produced by these two infection types following a non-human (rat) blood meal. When the *Wolbachia*-infected *Ae. aegypti* colonies were fed on rat blood but supplemented with additional amino acids, fecundity increased by approximately 15-20 eggs and egg hatch rate increased by 30-40 %. This suggests that a rescue effect may be possible if selection to the alternative blood source cannot be achieved to reach the phenotypic output observed on human blood. Groups that are seeking to mass rear large colony numbers of *Ae. aegypti* may still be able to produce locally competitive mosquitoes with non-human blood if a supplementation of key nutrients such as amino acids is supplied.

With regards to the consequences for large-scale *Wolbachia* mediated arboviral control strategies, there are two methods using *Wolbachia-*infected *Ae. aegypti* currently in development. The first method under investigation involves population suppression induced by releasing mass quantities of incompatible (*Wolbachia-*infected) males into uninfected populations. The *Wolbachia-*infected males mate with uninfected females, rendering the offspring inviable due to cytoplasmic incompatibility [7,39,40]. We have not tested the effects of alternate blood sources on incompatibility in our experiments, but incompatibility associated with *w*AlbB remains strong when males come from colonies that have been maintained on other blood sources (Nazni, pers. comm.). The second involves replacement of *Ae. aegypti* populations with *Wolbachia-*infected mosquitoes which then interferes with arboviral replication and transmission [41]. If large colonies of *Wolbachia*-infected *Ae. aegypti* are being raised in laboratory conditions using non-human blood sources, this may lead to the establishment of *Ae. aegypti* colonies intended for release with reduced and sub-optimal *Wolbachia* density (reduced density relative to density levels achieved by human blood meals). When these colonies are released into field settings, the ability of the invading *Wolbachia*-infected *Ae. aegypti* to replace naïve target populations will be diminished [42]. This will likely hinder *Wolbachia* invasion and replacement strategies. A decrease in *Wolbachia* density could also influence the ability of *Wolbachia* to block disease transmission which seems to relate at least partly to *Wolbachia* density in the host [18]. Should invasion succeed, the ability of *Wolbachia-*infected *Ae. aegypti* to supress arboviral transmission may still be insufficient, but only if low *Wolbachia* densities are inherited.

## Author Contributions

Conceptualization, A.A.H., P.A.R.; Methodology, V.P., E.C., P.A.R., J.K.A. and A.A.H.; Formal Analysis, A.A.H., V.P., and E.C.; Investigation, V.P and E.C.; Writing—Original Draft Preparation, V.P., E.C., and A.A.H.; Writing—Review and Editing, V.P., E.C., P.A.R., J.K.A. and A.A.H.; Visualization, V.P.; Supervision, A.A.H.; Funding Acquisition, A.A.H.

## Acknowledgments

The authors thank Nancy M. Endersby-Harshman and Ashley G. Callahan for technical assistance and valuable discussions.

